# Hummingbird abundance is related to food resources availability in a temperate forest of central Mexico

**DOI:** 10.1101/2021.10.30.466620

**Authors:** Mario Abraham Vazquez-Buitrón, Miguel Angel Salinas-Melgoza, Vicente Salinas-Melgoza, Alejandro Salinas-Melgoza

## Abstract

One strategy animals perform to cope scarcity of food resources is to shift in number of individuals according towards areas with available resources. This strategy can be more marked in species that are constrained by high energetic requirements such as hummingbirds. We aim to determine the extent to which the availability of food resources could be predictor of hummingbird numbers within and across hummingbird species in a temperate forest of central Mexico. We anticipate fluctuations in the number of hummingbirds grouped by species is best explained by monthly fluctuations in flowering resources species compared to pooled data. Our results indicate all seven hummingbird species fluctuate monthly in number across the year, which corresponds to monthly fluctuations of plant species they feed upon. The *Basilinna leucotis* and the *Salvia elegans* were present and interact in the study area almost all year-round, guiding the abundance pattern of both plants and hummingbirds in the study site. Generalized Linear Mixed Models indicate that although considering the abundance of the number of flowers for all plant species together could explain the fluctuation in all hummingbird species pooled together, considering both plant species and hummingbird species separately can provide a better explanation for changes in bird abundance. The model that analyzed species indicate that the interaction between the year-round species *B. leucotis* and *S. elegans* recorded the highest significant size effect. Our results highlight the fact that abundant species guiding abundance patterns could obscure by-species hummingbird trends and the processes guiding their patterns of abundance. We point out the need for performing adequate analytical approaches that can detect important biological interactions, and the likelihood of changes in habitat changing the pattern observed.

**RESUMEN:** Una estrategia los colibríes usan para lidiar con la escasez de alimento es realizar cambios en número de individuos a zonas con recursos disponibles. Esta estrategia es más marcada en especies que son limitadas por altos requerimientos energéticos como los colibríes. Nuestro objetivo fue determinar el grado con el cual la disponibilidad de alimento puede ser un predictor de número de individuos en un bosque templado del Centro de México. Anticipamos fluctuaciones en número de colibríes agrupados por especies serían mejor explicadas por fluctuaciones mensuales en recursos florales al comparar con datos combinados. Nuestros resultados indican que las siete especies de colibríes fluctuaron mensualmente en números a lo largo del año de acuerdo a fluctuaciones mensuales de las especies de plantas que ellos se alimentan. *Basilinna leucotis* y *Salvia elegans* estuvieron presente e interactuaron en la zona de estudio casi todo el año, guiando el patrón de abundancia tanto de colibríes como de plantas. Los Modelos Lineales Generalizados Mixtos indicaron que aunque considerar la abundancia del número de flores para todas las especies juntas podría explicar la fluctuación en todas las especies juntas, considerar las especies de colibríes y de plantas por separado provee una mejor explicación para los cambios en abundancia de aves. El modelo analizando especies indicó que la interacción entre las especies anuales *B. leucotis* y *S. elegans* tuvieron el tamaño del efecto significativo más alto. Nuestros resultados destacan el hecho de que especies abundantes guiando el patrón de abundancia podrían obscurecer tendencias especificas por especie y los procesos guiando su patrón de abundancia. Indicamos la necesidad de usar enfoques analíticos adecuados que puedan detectar interacciones biológicas importantes, así como la probabilidad de que cambios en el hábitat pueden cambiar el patrón observado.

*Palabras clave:* Bosque templado, abundancia estacional, uso del hábitat, asociación colibrí-comida, fluctuaciones temporales en abundancia

*Lay summary:*

- Hummingbirds, as many animals with high energy requirements, might cope with food resources shortage using different strategies
- One strategy to face food shortage is the local shifting in number of individuals.
- We used monthly surveys of hummingbirds and flowering plants in a temperate forest of central Mexico to evaluate the association of resources availability, seasonality, species identity, and vegetation condition on hummingbird abundance.
- All seven hummingbird species fluctuate in number across the year, which matches to fluctuations of plant species they feed upon.
- Hummingbird species *Basilinna leucotis* and the plant species *Salvia elegans* are the most abundant and largely guide the general abundance pattern.
- Both plant and hummingbird species separately are better explanting changes in hummingbirds’ abundance than species abundance combined.
- Abundant species may guide the plant and hummingbird abundance patterns, which complicates understanding underlying processes per species for the whole community.
- Given the current trends of habitats modification and the fact that habitat condition may influence the presence of key plant species for hummingbirds, we need to protect habitats where these key food plant species for hummingbirds are, particularly if they are specific to habitats

## INTRODUCTION

Understanding patterns of species abundance and distribution and the factors that govern them is a key topic in ecology. The abundance and distribution of animals can fluctuate across time and space. Resident species are animal species that show a more consistent pattern in their occurrence in a given space with moderate or no fluctuations in numbers. Other animal species show contrastingly fluctuating patterns. For example, a species may not occupy a given region at a given time but may increase their abundance at some point in the future (Brown et al. 1995). These fluctuations in abundance and distribution of individuals can impact the regulation of interspecific interactions (Souza et al. 2018), population dynamics (Jones et al. 2003), and community structuring (Holmes et al. 1986, Ostfeld and Keesing 2000) via a number of factors including food availability, habitat features, and environmental variables such as temperature or rainfall (Holmes et al. 1986, Levey and Stiles 1992, Brown et al. 1995, Ostfeld and Keesing 2000, White 2008, Mutshinda et al. 2009). The main factor driving the abundance of individuals is the availability of food resources (Wolf et al. 1976). Food scarcity especially can drive fluctuations in abundance via energetic constraints, particularly in species with intrinsic characteristics that impose high energetic demands.

### Ubiquity of Food Resources Scarcity in Hummingbirds

Hummingbirds have high energetic requirements (Powers and Conley 1994, Gass et al. 1999), which makes them more likely to be constrained by food availability. This means hummingbirds are highly dependent on a constant supply of highly energetic food resources such as flower nectar to fulfill those energetic requirements. Dependency on a continuous supply of food makes both temporal and spatial fluctuations in plant food resources provided by plants species important drivers of the pattern of hummingbird abundance.

Food resource availability for hummingbirds can fluctuate at different spatial and temporal scales due to several factors. Plant species that serve as food resources for hummingbirds may show contrasting temporal patterns of food availability. For example, this can result in constant availability of food resource with peaks of high abundance of flowering resources that provide abundant food resources. Conversely, other plant species occur temporally staggered between peaks of food availability which can serve as ephemeral sources of food supply during critical times of food shortage (Wolf et al. 1976, Kuban and Neill 1980, Stiles 1980, Stiles 1985, Arizmendi and Ornelas 1990, Poulin, et al. 1993, Gutiérrez-Zamora et al. 2004, Lara 2006, Cotton 2007, López-Segoviano 2018). Fluctuations can also occur spatially as habitat type can be a key driver shaping the localized spatial pattern of occurrence and abundance of flowering resources used by hummingbirds (Wolf et al. 1976, Kuban and Neill 1980, Stiles 1980, 1985, Arizmendi and Ornelas 1990, Wolf and Hainsworth 1990, Lara 2006, Cotton 2007). Fluctuation in food resources constrained by habitat type can also work in tandem with seasonality, as plant species flowering can be associated with either forested or open vegetation during specific times of the year (Kuban and Neill 1980, Stouffer and Bierregaard 1995, Arizmendi 2001, Lara 2006). Habitat modification can also modify the conditions where plant species originally occur, in turn, leading to changes in food availability (Stiles 1980, Chaturvedi et al. 2017; Sanaphre-Villanueva et al., 2017; Zubair et al. 2016).

This temporal and spatial fluctuation can lead to an asymmetrical temporal and spatially distribution of flowering resources, with some plant species found preferentially occurring in some habitat types in a specific season, and some other times and locations with low or no resources available. Taken together, this may lead to the impression of an increased degree of uncertainty in food availability for hummingbirds; however, the constant turnover of plants species due to the staggered flowering brings relief during times of scarcity. Despite this, hummingbird morphological constraints and specific species’ diet preference (Maglianesi et al. 2014) may preclude certain hummingbird species from exploiting available resources, causing individuals to experience a food shortage at some point during the year.

### Strategies Used by Hummingbirds to Cope with Fluctuations in Food Abundance

As a result of their dependence in highly energetic food resources, hummingbirds show a number of strategies to cope with fluctuations in food resources and food scarcity which help maximize foraging efficiency. Behavioral strategies include increasing the efficiency of nectar extraction (Wolf et al. 1976) or reducing intervals between feeding bouts and adjusting their foraging intervals according to nectar-renewal rates (González-Gómez et al. 2011). Hummingbirds also remember flowers with low nectar payoff during foraging trips and avoid visiting empty flowers in the future (Healy and Hurly 1995). It is suggested that plant species utilized by hummingbirds produce the most nectar (Feinsinger 1976); however, hummingbird preference for plant species may not be related to the nectar amount they produce (Arizmendi 2001).

Hummingbirds also use strategies that include shift in numbers due to changes in food resources availability. Some studies recorded the occurrence of reciprocal patterns of abundance at regional scales (Reviewed in: Fleming et al. 2005). While some sites report a reduction in the number of hummingbirds, other sites record increases; however, no association with food resources are performed. This suggests fluctuations in flower availability can have a strong influence in hummingbird abundance and richness. This suggests hummingbirds track actively their food resources (Cotton 2007) performing movements at different scales (Stiles 1980, Stouffer and Bierregaard 1995, Arizmendi 2001, Lara 2006, Cotton 2007), and has been confirmed from marked individuals (Stiles 1980). Other studies have evaluated this relationship, which can lead to higher or lower local abundances of hummingbirds. The number of individuals can increase or decreases steadily accordingly with either food resource availability or energy produced by plant species to cope with constant fluctuations in food resources, which increase the matching of hummingbird abundance with that of flowering resources (Feinsinger 1976, Montgomerie and Gass 1981, Stiles 1985, Lara 2006, Cotton 2007, López-Segoviano et al. 2018). Hummingbirds can also show an abrupt local increase in numbers, a strategy normally associated with latitudinal migrants during their movement to wintering grounds (Kuban and Neill 1980, Mckinney et al. 2012). Species such as *Selasphorus platycercus* and *Selasphorus rufus* suddenly increase numbers at stopovers during peak flowering time periods by matching the timing of occurrence at stopovers and the timing of flowering phenology of preferred plant species.

### Improving Our Understanding of Food Availability-hummingbird Abundance Association

The way that the association between hummingbird abundances and their food resources has been evaluated includes several analytical approaches which has implications for our understanding of this association. While some studies pooled community abundance of hummingbird species (Feinsinger 1976, Montgomerie and Gass 1981, Stiles 1985, Cotton 2007, Abrahamczyk and Kessler 2010, Bustamante-Castillo 2018), others separated abundance of individuals by species (Feinsinger 1976, Stiles 1985, Bustamante-Castillo 2018). Pooling abundance could obscure by-species trends and the relationships guiding their patterns of abundance due to several reasons. First, hummingbirds and the plant species they visit can show morphological matching which increases the tendency of non-random visitation patterns (Sonne et al. 2018, Rodríguez-Flores et al. 2019, Sonne et al. 2020). Second, the temporal or seasonal pattern of plant occurrence affects the frequency of plant-hummingbird interactions which can be skewed by hummingbird migratory status. Third, some plant species may not be abundant, but are important food resources for hummingbird species in critical times of the year. For example, *Hamelia patens* flowers produce a large amount of nectar making it a highly preferred resource to hummingbirds; however, they are not an abundant species (Feinsinger 1976). Fourth, hummingbird species abundance can bias the community patterns of abundance when species abundances are pooled (Arizmendi and Ornelas, 1990, Bustamante-Castillo et al. 2018). In this scenario, an overabundance of a hummingbird or plant species may overemphasize their importance at a community-level. An association in this scenario could indicate the influence of fluctuations in food resources used by the abundant hummingbird species rather than for all species.

In previous studies, not all evaluations are clear on what approach could be better to help understanding hummingbird response or the implications of pooling data together. The concurrent evaluation of both pooled abundances and separately can improve our understanding of these associations, considering abundances separately can not only help disentangling those associations by plant species, but also provide better explanations to patterns. Examples of this are *Phaethornis guy* and *Philodice bryantae*, whose abundances were found to be associated with the number of all flowers; however, the association was stronger when just the flowers of *Heliconia* community and *Inga brenessi* were considered respectively for each species of hummingbird (Feinsinger 1976, Stiles 1985). Considering the abundances of the different species pooled to evaluate food and hummingbird abundances association may obscure important biological interactions due to an inadequate scale of analysis.

We evaluate the influence of food resources fluctuation in the abundance of hummingbirds in a temperate system of Central Mexico high plateau. Anthropogenic activities in this study area have opened the forest and introduced exotic plants introduced by human activities that can be alternative food resources for hummingbirds. The introduction of exotic plants can influence the pattern of abundance of hummingbirds via changes in plant species hummingbirds feed upon. We hypothesize the increase in hummingbirds’ numbers is due to an increase in the availability of their main dietary flowering resource. We anticipate these associations would be also influenced by the degree of modification of the forest and by seasonality.

## METHODS

### Study Site

We performed this study at the Área Natural Protegida La Sierra de Los Agustinos Reserve in Guanajuato State, Mexico (20° 13’North and 100° 40’ West). This protected area is located at the Sierra de Los Agustinos mountain range at the central Mexico Highlands. This reserve is 19, 246.00 hectares (CONABIO 2012) and comprise three municipalities: Acámbaro, Jerécuaro and Tarimoro. The highest altitude is 3, 110 masl, where a temperate forest is the dominant type of vegetation. This mountain range is surrounded by agriculture areas at its base. The rainy season runs from June to October, and the dry season runs from November to May. Peak rainfall occurs between July and August when average monthly rainfall is about 100 mm.

#### Vegetation modification

The area has five distinguishable vegetation types that are somewhat stratified by altitude but can overlap. The pine forest, oak-pine forest, forest stands are dominated by oak, subtropical vegetation, and grasslands (CONABIO 2012; Rzedowski and Rzedowski 2005). Although the area where the study was conducted is now a protected area, the temperate forest was modified for several years in the past. Two type of vegetation conditions can be distinguished based on habitat modification. The first condition is closed vegetation, composed of denser vegetation mostly dominated by trees from genus *Pinus* spp. and *Quercus* spp.. The second condition is open canopy vegetation, with a contrasting high degree of modification, where original trees are scarce and herbs dominate (CONABIO 2012).

### Study Design

We performed monthly sampling along 1 km long-10 m wide transects in the study area. To obtain a representative sample of local conditions, six transects were distributed, three in each forested and open vegetation. Surveys were performed monthly from March 2018 through February 2019, except for October 2018. Each transect was surveyed one day each month. Transects were at least 500 meters apart to reduce the likelihood of counting the same hummingbird individual more than once during surveys. We performed estimations of hummingbird abundances and food availability along these transects.

#### Estimation of hummingbird abundance and food availability

We obtained the number of hummingbirds present monthly in two ways. First, we surveyed 25 meter fixed-radius point counts. We identified number of individuals per species during 10 minute period observations at each point count. Second, we used the number of birds per species captured in mist nests. Hummingbirds were identified at the species level with binoculars and the identification guide of Colibríes de México y Norteamérica (Arizmendi y Berlanga 2014). Hummingbird abundance was estimated as the total number of individuals per species at each transect in each monthly survey.

We estimated the abundance of food resources by performing surveys of flowers along transects at the same time when hummingbird numbers were obtained. The total number of flowers per plant species counted at each transect was used as a proxy for food availability. We also obtained pollen grain samples from these plants to generate a reference pollen library that we used to identify pollen grains found in hummingbird pollen loads. Although we evaluated the abundance of flowers for the whole community of flowering plants, we report abundance only for those top seven ranked plant species. Values are indicated as averages and standard deviations.

#### Evaluation of plant species used by hummingbirds

We performed an evaluation of the plant species used by hummingbirds combining two approaches. First, we placed five mist nets at transects where hummingbirds’ abundance was obtained during each survey. Mist nets were placed where more flowers were available in each transect to increase the likelihood of capturing hummingbirds with pollen. Mist nets were opened from 7:00 am to 2:00 pm. Pollen load in combination with the pollen library was used to obtain information on the flowers visited by hummingbirds. Pollen load was estimated from pollen sample obtained from the hummingbird’s bill, crown feathers, and chin and throat feathers using a gelatin dyed with fucsina. Pollen samples were observed in an optic microscope at 40X. We combined pollen records by plant species from pollen loads and flower visitation observations as an indication of the relative contribution of each plant species to hummingbirds’ diet. Second, we obtained observations of hummingbirds visiting flowers along transects, at count points, and outside sampling areas. These individuals were not captured, but both hummingbird species and the food resource species were identified and recorded, as well as the surrounding vegetation where the observation occurred. We defined the six top-ranked plant species as the main food items in the hummingbird community by their relative contribution to diet and by also the timing of flowering compared to other plant species. Some plant species only produced flowers during a specific time of the year, and they were important plant species at that time of the year.

### Statistical Analyses

We used Generalized Linear Mixed Models (GLMM) with a log link to tackle the lack of independence in repeated measurements. The dataset with monthly flower abundance included the abundance for the six key plant species for hummingbirds in the study area: *Salvia elegans, Penstemon roseus, Macromeria longiflora, Opuntia* spp. *Leonotis nepetifolia*, and *Agave* spp. First, we first constructed a model with pooled species abundance per transect per month as the response variable. Vegetation condition, season, and standardized number of flowers for all plant species pooled together per transect per month were considered fixed effects. We investigated whether these variables could explain hummingbird variance for all species pooled per month. This would enable us to make comparisons with other studies that compare pooled flower abundances and indicate hummingbirds’ response to fluctuations in flowers as a whole. Both condition and seasonality had two levels: closed vegetation and open vegetation, and the rainy and the dry season respectively.

A second non-zero-inflated Generalized Linear Mixed Model (GLMM) was constructed. Here, vegetation condition, season, hummingbird species identity as a dummy variable, and standardized number of individuals per plant species were included as fixed effects. The abundance for each hummingbird species separately was considered as response variable. From pollen load alone, it was not possible to identify the key plant species to the species level the pollen type of one of, thus it was not possible to relate it to an estimation of availability. Hence, unidentified key pollen plant morphs were removed from further statistical analyses. Analyzing the six key plant species and all hummingbird species separately helped us to disentangle the effect of each flowering plant species and the individual response of each hummingbird species. This enabled us to determine if independent plant species contribute differentially to hummingbirds’ abundance pattern. Transects were considered a random effect across the sampling period for both models. We evaluated the dyadic interaction between the main hummingbird and plant species (*B. leucotis* and *S. elegans*), as they were the most abundant and more consistently present and interacting throughout the year. The interaction between *B. leucotis* with season and vegetation type was also evaluated.

We excluded *A. beryllina* from the abundance datasets because the species was recorded only once, with one individual during the study period. We determined the appropriate error distribution for response variables (Faraway 2016). A negative binomial type 2 error distribution was used for both models. We performed likelihood ratio test by comparing the full model including the effect of the fixed effect against the reduced model without the effect using a forward stepwise method. We used the *glmmTMB* package 1.0.2.1 (Brooks et al. 2017) for R 3.3.2 (R Development Core Team 2021) to perform GLMMs. We considered Marginal R^2^ as a measure of how well each model could help understanding response variables. Results are presented as means and standard deviations.

## RESULTS

### Food Resources Abundance

We recorded 16 plant species from sightings in transects surveys and from pollen load: *S. elegans, Castilleja tenuiflora, Lobelia laxiflora, P. roseus, Bouvardia longiflora, Opuntia spp., L. nepetifolia, Echeveria secunda, Loeselia mexicana, Salvia mexicana, M. longiflora, Salvia patens, Agave spp., Oenothera hartwegii, Scutellaria dumetorum, Ipomoea murucoides*. We also recorded pollen grains from two plant species for which it was not possible to assign a species identity.

The presence of all available flowering plant species and its abundance fluctuated monthly (Fig. 1). The lowest average of total number of flowers for all species combined per transect was recorded in July 2018, followed by June 2018. There were three peaks of flower production. March and August 2018 and February 2019. *S. elegans, L. nepetifolia, Opuntia spp*., and *P. roseus* were the most abundant plant species. The dry season recorded the largest percentage of flowers available for all plant species (76%; 1,436.5 ± 1153.0 average flowers per transect per month, Range = 411.8 – 3,228.8) in comparison with the rainy season (16%, 362.9 ± 359.9 average flowers per transect per month, Range = 123.2 - 398).

**Figure 1.**
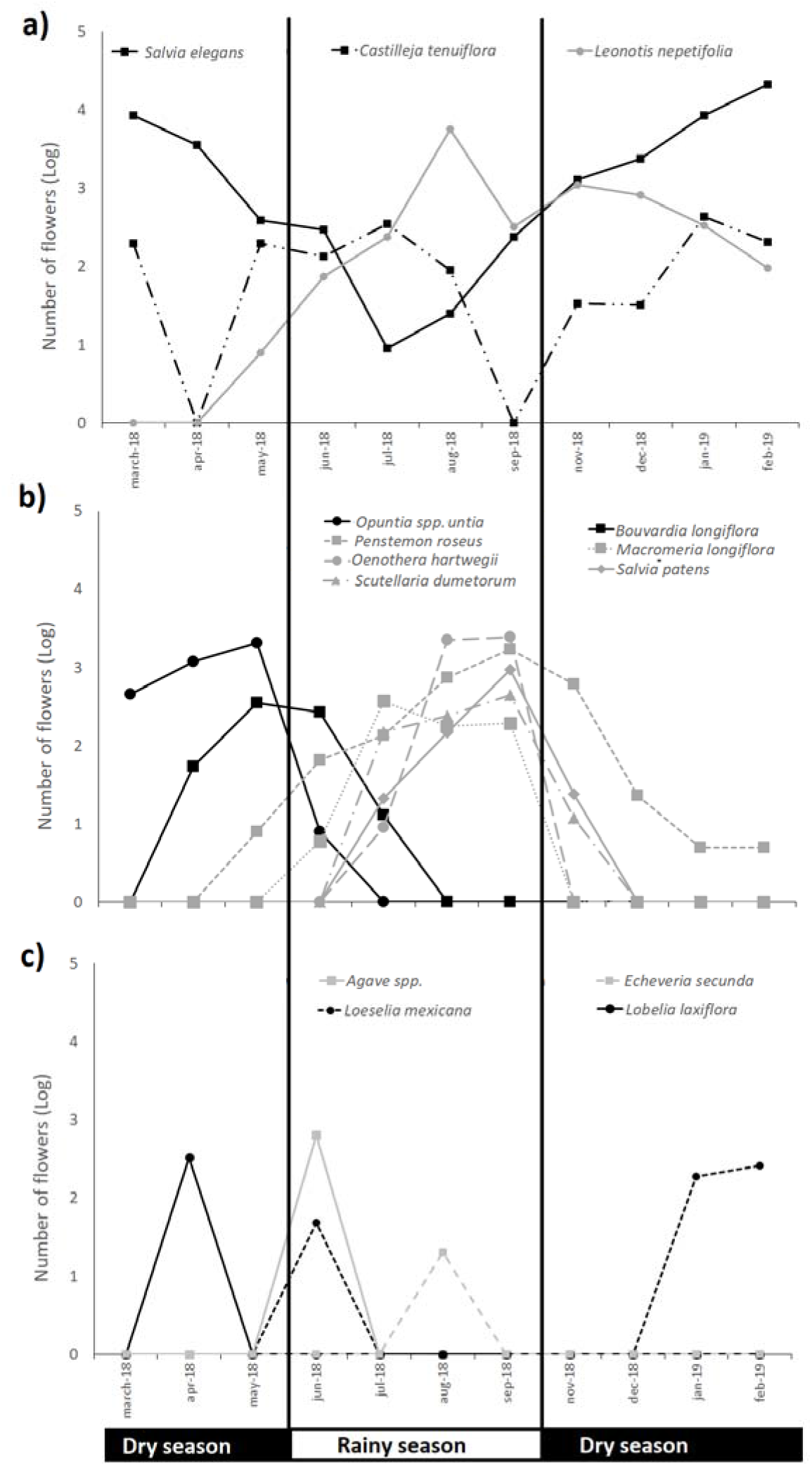
Abundance of key flowers available per transect per month for hummingbirds in the Sierra Los Agustinos Reserve during March 2018 through February 2019. a) Year-round plant species. b) Plant species predominantly flowering during the dry season (grey line) and plant species predominantly flowering during the rainy season (black line). c) Plant species flowering sporadically.

Turnover of available flowering plant species during the year, resulted in four temporal flowering strategies creating a seasonal fluctuation of available flowering resources: a)Plant species flowering practically throughout year, such as *S. elegans*, b) plant species flowering exclusively during the dry season to the beginning of the rainy season (Fig. 1b), c) plant species which produced flowers mostly during the rainy season, July-September, and d) plant species providing ephemeral flowering food for hummingbirds. *S. elegans* drove the flowering pattern (Fig. 1a), recording up to 68,271 flowers in total through the complete sampling period. *L. nepetifolia* was another species flowering entirely throughout the study period (May 2018 - March 2019) (Fig. 1a). Dry season flowering plants included *Opuntia* spp., which was recorded exclusively flowering from March-June, utilizes this strategy. March 2018 recorded the lowest number of plant species flowering (3 species), being *S. elegans* the most abundant in this month. Rainy season flowering plants included plant species such as *O. hartwegii, P. roseus, S. patens, S. dumetorum, M. longiflora, L. mexicana, L. laxiflora, E. secunda*, which produced flowers mostly through July-September (Fig 1b). The largest number of different plant species flowering occurred in August 2018 (10 plant species). Plant species with ephemeral flowering food resources included *Agave* spp., *I. murucoides, B. longiflora, L. Mexicana, L. laxiflora* and *E. secunda* (Fig. 1c). The flowering timing for these plant species was scattered throughout the year. Taken together, this indicates there is a marked fluctuation in food resources with an associated food scarcity occurring in June-July and by-species flower production is highly concentrated in a small-time window in February-March, driven by *S. elegans* due to its high abundance.

The availability of flowering resources per plant species had a pattern of occurrence associated with habitat condition. *S. elegans, C. tenuiflora, L. laxiflora, P. roseus, E. secunda, S. mexicana, M. longiflora, S. patens*, and *S. dumetorum* occurred preferentially in closed vegetation. While *B. longiflora, Opuntia* spp., *L. nepetifolia, L. mexicana, Agave* spp., *O. hartwegii, I. murucoides* occurred in open vegetation. *Opuntia* spp. was recorded exclusively during the dry season, while *Agave* spp. and *M. longiflora,* produced flowers exclusively in the rainy season (Fig. 1). Both *L. nepetifolia* and *P. roseus* were recorded flowering preferentially during the rainy season. *S. elegans* was the only plant species recorded preferentially during the dry season. This indicated clear by-species patterns of temporal and spatial occurrence with both *M. longiflora* and *P. roseus* exclusively producing flowers in the closed vegetation during the rainy season. While *L. nepetifolia* was recorded producing flowers in open vegetation during the rainy season; *S. elegans* showed an inverse pattern mostly producing flowers in the closed vegetation during the dry season.

### Hummingbird Abundance

We recorded seven hummingbird species in our study site through the sampling period: *Basilinna leucotis, Eugenes fulgens, Leucolia violiceps, S. rufus, Amazilia beryllina, Colibri thalassinus* and *Cynanthus latirostris*. The highest number of species was recorded in both July 2018 and September 2018 (Fig. 2). April 2018 recorded the lowest number of species, when only the year-round *B. leucotis* occurred. The hummingbird community was dominated by the year-round resident species *B. leucotis* (Figure 2a), which recorded its peak in February 2019.The lowest number of *B. leucotis* was recorded in May 2018 and August 2018. *E. fulgens* was the second most common species (Fig. 2a). Seasonal fluctuations were also observed. The year-round species *B. leucotis* dominated the community during the dry season. While *E. fulgens, L. violiceps, C. latirostris, C. thalassinus*, and *S. rufus* abundance was larger in the rainy season (Fig.2).

**Figure 2.**
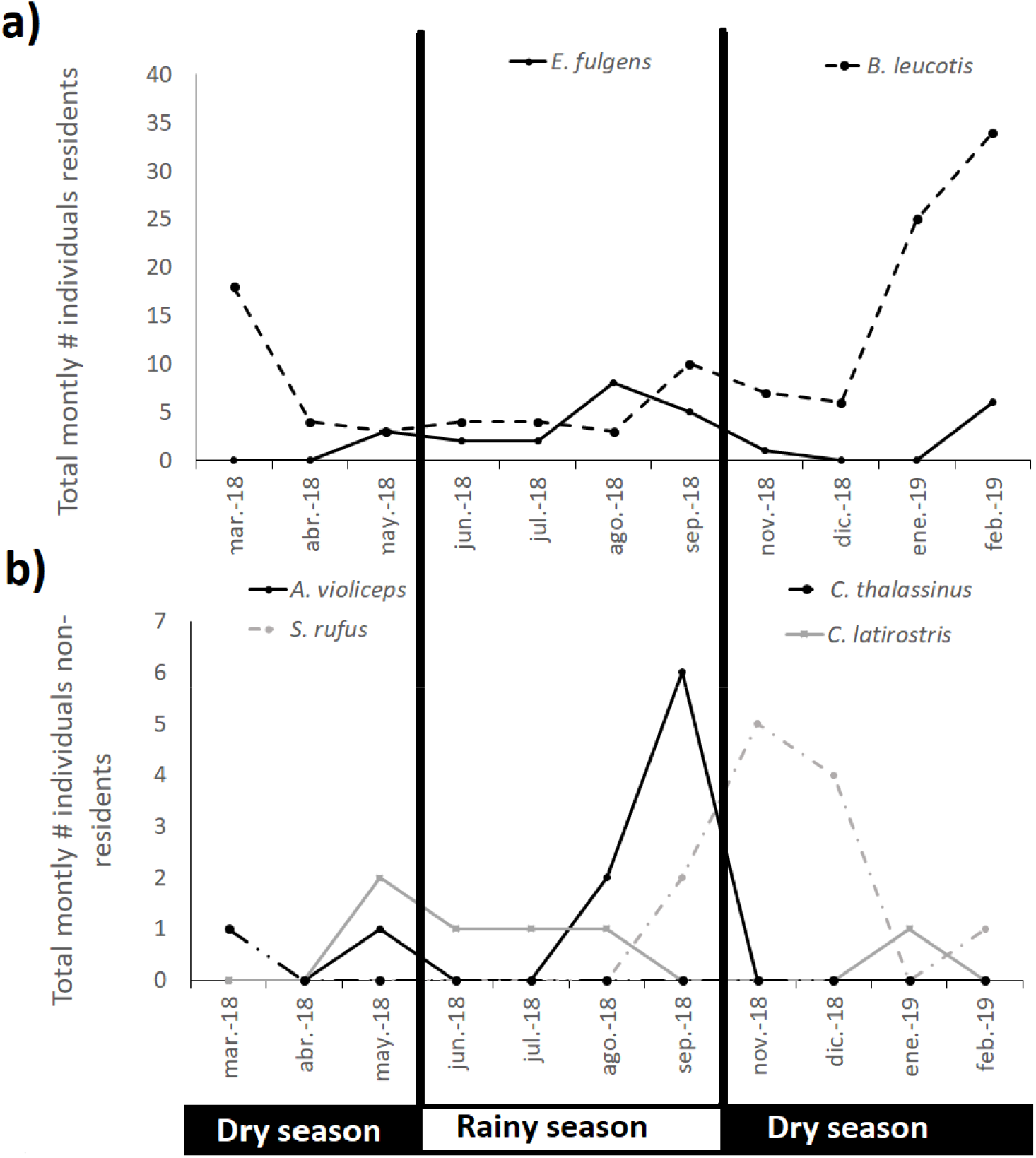
Abundance of hummingbird species per transect per month in the Sierra Los Agustinos Reserve during March 2018 through February 2019. a) Year-round hummingbird species. b) Non-year-round hummingbird species.

The pattern of hummingbird occurrence also varied by vegetation condition as species showed a high segregation by vegetation type. *A. beryllina, L. violiceps*, and *C. latirostris* were only recorded in open vegetation. *C. thalassinus*, *S. rufus*, and *E. fulgens* were recorded preferentially in open vegetation; however, *S. rufus* and *E. fulgens* could not be assigned to a vegetation condition conclusively. The most abundant hummingbird species in our community, *B. leucotis*, preferred closed vegetation as 90% of individuals were recorded in this condition. Most of the records for *L. violiceps* and *E. fulgens* occurred during the rainy season (Fig. 2). While *B. leucotis* recorded larger numbers in the dry season, *S. rufus* recorded more individual at the end of the rainy season-beginning of the dry season. *C. latirostris* was present in both seasons, although preferentially in the rainy season. *A. beryllina* was exclusively recorded during the rainy season. This indicated hummingbird species showed a temporal and spatial segregation, with *C. thalassinus*, *L. violiceps*, and *E. fulgens* mostly occurring during in the open vegetation during the rainy season. *S. rufus* occurring mostly in the open vegetation during the dry season. *B. leucotis* heavily occurred in the closed vegetation during the dry season. While *C. latirostris* occurred mainly in the open vegetation, its seasonal occurrence did not show a clear tendency.

### Pattern of Food Items Use by Hummingbird Species

Pollen loads collected from the entire hummingbird community recorded 19 plant species. We identified 14 food items to the species level: *S. elegans, L. nepetifolia, M. longiflora, P. roseus, S. mexicana, S. patens, L. mexicana, C. tenuiflora, O. hartwegii, S. dumetorum, L. laxiflora, B. longiflora, E. secunda*, and *I. murucoides*. We identified two food items at the genus level: *Opuntia* spp., and *Agave* spp., but we were not able to identify three food items at the genus level. The six important key food items that were possible to assign a species and genus level accounted for 76% of all food items recorded. The use of the year-round plant species *S. elegans* was monopolized by *B. leucotis. L. nepetifolia* was mostly used by *L. violiceps*, followed by *S. rufus*. Half the records of *M. longiflora* corresponded to use by *E. fulgens*, followed by *L. violiceps. E. fulgens* and *B. leucotis* recorded almost a similar use of *P. roseus*. The main plant species recorded in pollen loads for all hummingbirds were *S. elegans, L. nepetifolia, M. longiflora. S. elegans* was the main food item for *C. thalassinus* (80%) and *B. leucotis* (80%). *L. nepetifolia* was the main food item recorded in *L. violiceps* (35%), *C. latirostris* (67%), and *S. rufus* (68%)*. M. longiflora* was the main food item in *E. fulgens* (35%). Plant species showing a narrow timing of occurrence by the end of the dry season such as *Opuntia* spp. and *Agave* spp. made up over a third of *L. violiceps* and *E. fulgens*.

GLMM indicated that all variables and their interactions explained a positive variation in this response variable (Table 1) when hummingbird abundance for all species was pooled per month. The largest effect resulted from the influence of all six plant species pooled followed by open vegetation. GLMM also indicated that hummingbirds’ abundance pattern could be explained for each species when species were considered separately, with the exception of *S. rufus*. The largest effect was due to the year-round hummingbird species *B. leucotis*, followed by *L. violiceps* (Table 2). Our results indicate that abundance of both the year-round hummingbird *B. leucotis* and its main food source, the year-round plant species *S. elegans*, are interacting. This suggests that abundance pattern observed for the whole hummingbird community could be driven by *B. leucotis* influenced by *S. elegans*. Comparisons of the Marginal R^2^ values for each model indicated analyzing hummingbird and plant species abundance independently outperformed the model when abundances were pooled. Thus, analyzing plant and hummingbird species separately may provide a better explanation for pattern of abundance, probably as a result of partitioning the relative variance contribution by species (Tables 1 and 2).

**Table 1.** Mod estimates from GLMM analyses on total number of all hummingbird individuals pooled for all species per month in the Sierra de Los Agustinos Protected Area. Estimates are indicated with confidence interval in parenthesis

| Predictors | Incidence Rate Ratios (CI) | *p* |
| --- | --- | --- |
| (Intercept) | 20.68 (17.39 – 24.59) | **<0.001** |
| Dry season | 0.64 (0.54 – 0.76) | **<0.001** |
| All six plant species pooled | 3.77 (2.64 – 5.38) | **<0.001** |
| All six plant species pooled : Open areas | 0.25 (0.17 – 0.36) | **<0.001** |
| All six plant species pooled: Dry season | 0.36 (0.26 – 0.52) | **<0.001** |
| Open vegetation | 0.81 (0.66 – 0.98) | **0.03** |
| **Random Effects** |  |  |
| $\sigma^2$ | 0.21 | |
| ICC | 0.02 |  |
| Marginal R<sup>2</sup> | 0.33 |  |

Rainy season, closed vegetation, and plant abundance for all six plant species interacting with closed vegetation and rainy season are included in the Intercept as references for inter-group comparisons.

**Table 2.** Model estimates from GLMM analyses on the abundance of each hummingbird species separately in the Sierra de Los Agustinos Protected Area. Estimates are indicated with confidence interval in parenthesis

| Predictors | Incidence Rate Ratios (CI) | *p* |
| --- | --- | --- |
| (Intercept) | 0.02 (0.01 – 0.06) | **<0.001** |
| *L. violiceps* | 75.92 (31.14 – 185.09) | **<0.001** |
| *C. latirostris* | 27.19 (9.60 – 77.03) | **<0.001** |
| *C. thalassinus* | 43.94 (11.50 – 167.93) | **<0.001** |
| Open vegetation | 1.80 (0.90 – 3.61) | 0.09 |
| *E. fulgens* | 48.38 (23.46 – 99.78) | **<0.001** |
| *B. leucotis* | 100.93 (36.79 – 276.90) | **<0.001** |
| *B. leucotis* : Open vegetation | 0.42 (0.16 – 1.10) | 0.08 |
| *B. leucotis* : *S. elegans* | 1.58 (1.35 – 1.85) | **<0.001** |
| *B. leucotis* : Dry season | 0.90 (0.39 – 2.09) | 0.81 |
| Dry season | 1.16 (0.64 – 2.11) | 0.62 |
| **Zero-Inflated Model** |  |  |
| (Intercept) | 0 (0.00 – Inf) | 0.99 |
| **Random Effects** |  |  |
| $\sigma^2$ | 1.5 | |
| Marginal R<sup>2</sup> | 0.62 |  |

Factors of rainy season, closed vegetation condition, and the interaction of both *B. leucotis* abundance with closed vegetation and with rainy season are included in the intercept as references for inter-group comparisons.

## DISCUSSION

Here we explored the shift in hummingbird abundance as a response by to availability of food resources with number of flowers as a proxy. We found an association of pooled hummingbird species abundance with factors of seasonality, vegetation condition, and flowering resource availability of for all six main food plant species pooled together. With the exception of *S. rufus*, species identity, rainy season, and vegetation condition also influenced hummingbird abundance pattern when we considered the abundance of each hummingbird and plant species separately. The interaction between *B. leucotis* and *S. elegans* also played a role in hummingbird abundance; however, *B. leucotis* played the strongest role. Our data highlight that fluctuations in food availability can be influenced by habitat and seasonality for both hummingbird communities as a whole and for particular species. Our data also highlight the positive steady response of this avian group to fluctuations in food resources which has been indicated in other studies and for a variety of animal species. Hummingbirds’ abundance can be constrained by the availability of food resources due to its high reliance on continuous energy supply. This study increases our understanding of the processes governing hummingbird abundances and different strategies this avian group can perform in a temperate forest to cope with food scarcity.

Here we used the same data in two ways to evaluate the response of hummingbird abundance to fluctuations in flowering resources. Although both approaches explain variation in hummingbird abundance, considering each species separately perform better at disentangling the influence of abundance drivers. Additionally, pooling data together could potentially obscure by-species patterns. Considering pooled abundance all hummingbird and plant species can give us insights for the whole community of hummingbirds, but it does not provide information on how particular species are responding to specific factors driving their abundance. For example, *S. elegans* was the most abundant and most consistent plant species flowering throughout most of the year, which could potentially drive the pattern of flowering for the whole community of plants flowering when pooling data. Thus, pooling data of *S. elegans* with abundance of other plant species could mask their effect on hummingbird abundance.

We also found plant species differ in the flowering timing, creating a plant species turnover. This suggests the community of hummingbirds have flowering resources staggered through the year that could provide a continuous supply of food resources (Stiles 1985). In addition, plants species eaten by hummingbirds such as *M. longiflora, Opuntia* spp. and *Agave* spp. have flowering peaks that occur infrequently throughout the year matching the lowest availability of food resources. *B. leucotis* and *C. thalasinus* preferred *S. elegans* in their diets, while *S. rufus, C. latirostris*, and *L. violiceps* preferentially included *L. nepetifolia* in their diets. Because these plant species are strongly included in the hummingbirds’ diet, they are considered to be key flower resources providing some relief in critical times of food supply. The latter hummingbird species could suffer from a food shortage in July due to the low availability of their preferred diet items, creating a by-plant specific dependence of species based on food preferences. Hummingbird species show a preference for a reduced number of plant species (Céspedes, et al. 2019, Simmons et al. 2019, Rodríguez-Flores et al. 2019), which makes them more sensitive to fluctuations in food resources despite potential plant species flowering. This reinforces the idea that an analysis using abundance for hummingbirds and plant species pooled is not able to disentangle the effect of preferences and would lead to spurious relationships driven by most abundant flowering species.

Here we suggest using analytical approaches that are adequate at meaningful ecological scales such as the hummingbird species or community level analysis. Particularly, we think the use of hummingbird and plant species abundances separately could increase our understanding of how hummingbirds’ response to drives, as it would indicate the relative influence of explanatory factors to fluctuations in hummingbird abundance.

When the data were analyzed separately we found that the two year-round species, *S. elegans* and *B. leucotis*, are in close temporal association and play a major role in driving hummingbird abundance. This is the only dyadic interaction between a hummingbird and its main flower resource that was associated in our data. First, *S. elegans* comprises 80% of the diet of *B. leucotis*, meaning this hummingbird species has a strong preference for this flowering resource. Number of individuals of year-round hummingbird species such as *Philodice bryantae, Eutoxeres aquila, Phaethornis guy*, and *Doryfera ludovicae* are associated with their main food resources (Feinsinger 1976, Stiles 1985), indicating that tracking the main food item in hummingbirds’ diets may give insights on the drivers of abundance pattern for specific species. Second, *S. elegans* and *B. leucotis* are the most abundant food resources and hummingbird species respectively in the study region. Thus, it is not surprising they are guiding both abundance patterns of this system. Third, *S. elegans* recorded the lowest availability of flowers resources from April through August and its peak occurred in February 2018, which matches the lowest and highest numbers of individuals for *B. leucotis*. Finally, although nectivorous species can regularly adjust their diet (Fleming et al. 1993), it is common that dominant hummingbird species preferentially use plant species that are more abundant or with higher sugar concentration (Lara 2006, López-Segoviano et al. 2018). We did not collect information on flowers’ energy content and the hierarchical system in this hummingbird community; however, it was possible to identify *B. leucotis* was the dominant species in the hierarchy system in our study site, keeping well delimited territories defended against even larger species such as *E. fulgens* (Vazquez-Buitron, M. A. personal observations).

The close association between these two year-round species could reflect the uncertainty of food resources supply for *B. leucotis* by relying on flower resources that are more continuously available throughout the year (Lara 2006). The high degree of diet preference of *B. leucotis* could also mean the species is not as plastic to fluctuations in its main food resources and is obligated to respond to fluctuations of *S. elegans*, regardless of its hierarchical status. Alternatively, the payoff of alternative flowering resources for *B. leucotis* during times of scarcity could be so low that the species does not fulfill its energetic requirements, leading to a local shift in numbers rather than showing plasticity in its diet.

Our model explained the pattern of abundance of *L. violiceps* and *C. latirostris*, but not for *S. rufus*. In addition, it was not possible to relate these hummingbird species abundances to a given plant species. *S. rufus* was recorded in lower numbers in our study site from September to February, while *L. violiceps* and *C. latirostris* occurred from May to September. Pollen loads from *L. violiceps* and *C. latirostris* indicate they relied mainly on *L. nepetifolia*, a plant species only found in one transect in discrete and uncommon patches in the open vegetation condition in our study area, and for which flowering peaks in August and November. Thus, indicating that hummingbird species are actively tracking clumped and rare floral patches of *L. nepetifolia*, as indicated by their occurrence in pollen loads.

The temporal overlap in the occurrence of animal species with the phenology of their resources is expected to be larger in latitudinal migrants compared to short distance migrants or resident species (Jones et al. 2003). Likewise, *L. violiceps* and *C. latirostris* could not shift in numbers to fluctuations in food resources abundance, but rather synchronize their occurrence with the timing of occurrence and highest abundance of food resources produced by *L. nepetifolia* in the study area. Agreement in the timing of both hummingbird occurrence and the peak of flowering resources has been observed in *Selasphorus platycercus, S. rufus*, and *S. sasin* at stopover sites (Kuban and Neil 1980, Russell et al. 1994, Lara 2006, Lara et al. 2009, López-Segoviano et al. 2018); however, this was not observed for *S. rufus* in our study.

Another strategy that could not be identified by our approach could occur by the altitudinal migrant *C. thalassinus*, as it is present just during the rainy season and appears scattered along the year in our study site. *C. thalassinus* exploits *S. elegans* and *P. roseus* flowers. Although the largest availability of food resources in our study site does not occur during this season when this hummingbird species occurs, it is the time of the year when the largest number of plant species are flowering and when the largest availability of *P. roseus* occurs and *S. elegans* is sufficient bloom. This would represent hummingbird species coinciding with high quality availability of two of the most representative plant species in its diet and with a large variety of flower resources, which could be convenient for migratory species when performing altitudinal movements from lower grounds.

An alternative explanation for *L. violiceps, C. latirostris* and *C. thalassinus* not being associated with any given plant species is that our study design precluded us from capturing the temporal resolution at which abundance fluctuations occur. *B. leucotis* is a year-round species, meaning we could track monthly changes in its availability. López-Segoviano et al. (2018) found the number of individuals for the latitudinal migrant *S. rufus* and the altitudinal migrant *A. beryllina* were associated with their main food resources, *Salvia iodantha* and *Cestrum thyrsoideum*. Sampling in this study was performed every 10 days in a temperate forest of northwestern Mexico, to maximize temporal resolution of hummingbird-plant interactions during the two-and-a-half-month study. Monthly surveys were an adequate sampling resolution for year-round species but might not be adequate for other species with shorter residency time in our study area such as *L. violiceps, C. latirostris* and *C. thalassinus*. Thus, it is important to consider meaningful temporal scales when analyzing the relationship between hummingbird abundances and fluctuations in food resources to better reflect the scale of interactions occurring between hummingbirds and their resources. This suggests that the hummingbirds in our study might use a combination of strategies according to the hummingbird species.

We found that season and habitat conditions influence hummingbird abundance patterns. This was related to patterns of co-occurrence of plant species with particular hummingbird species. *S. elegans* was the only plant species recorded flowering almost exclusively during the dry season in closed vegetation, where it exclusively occurs (Lara 2006). As *S. elegans* was the most abundant flowering resource in our study site during the dry season which creates an asymmetrical temporal and spatial distribution of flowering resources, with the closed vegetation during the dry season providing most of the flowering resources exploited by several species. The asymmetric occurrence of flowering resources results in the largest amount of hummingbird individuals recording during the dry season, where the larger occurrence of both flowers and number of hummingbirds regularly occurs (Wolf et al. 1976, Stiles 1985, Bustamante-Castillo et al 2018). It is possible, however, to record resources and hummingbirds in larger numbers during the rainy season (Lara 2006).

Hummingbird species show a varying pattern of temporal and spatial overlap of occurrence with their main food resources. The ubiquity of *S. elegans* and its main role in *B. leucotis’* diet explains the high degree of temporal and spatial overlap by habitat type between them. Species such as *C. latirostris* and *L. violiceps* also had a close temporal and spatial overlap with their main food resource, *L. nepetifolia*, as they occurred in the open vegetation during the rainy season. These data highlight that plant species may be the main drivers of the hummingbird species temporal and spatial abundance by habitat type. In addition, *C. thalasinus* and *E. fulgens* co-occurred temporally with their food resources, *P. roseus* and *M. longiflora*, during the rainy season; however, they did not coincide spatially as resources were mostly in the closed vegetation. *C. thalasinus* represents an extreme on a spectrum of temporal and spatial agreement because it shows a contrasting pattern of occurrence with its main food resource, *S. elegans*. Hummingbirds preferentially occur in open vegetation during the rainy season and plant individuals occur preferentially in closed vegetation during the dry season. Because *S. elegans* comprises a large percentages of *C. thalasinus*’s diet suggests *C. thalasinus* could be tracking the occurrence of this year-round plant species, even in areas where this plant species occurs in low numbers. This could indicate these hummingbird species are performing local movements among patches of closed forested and open nonforested vegetation to track those few preferred diet items available in the area.

The temporal availability in food resources is probably the hardest constraint in food availability hummingbirds must face due to its reliability on constant energy supply. Our study shows hummingbirds may show a battery of strategies to cope with the food shortage that occurs in this temperate system in central Mexico. The clearest evidence is indicated by the association of two year-round species, *S. elegans* and *B leucotis*. Our data indicate hummingbirds perform a shift in numbers to response to fluctuations in flowering resources and highlight the importance of performing analyses at more biologically and temporally meaningful scales.

Our data indicate that some hummingbird species respond to changes in food availability and taken with the different patterns of resource availability between vegetation types, we emphasize the need to stop the reduction of closed, forested areas due to habitat modification. These conservation activities are necessary to protect hummingbird habitat. While migratory species track key plant species at stopover sites, yearlong-resident species rely on food resources occurring exclusively in closed vegetation. This makes yearlong-residents more susceptible to habitat modification. Plant species hummingbirds preferentially include in their core diet occurs in closed vegetation (Kuban and Neill 1980, Stouffer and Bierregaard 1995, Arizmendi 2001, Lara 2006), so vanishing closed vegetation would mean hummingbirds suffer due to an exacerbated food shortage in harsh times. Flower resources, such as *Agave* spp. and *Opuntia* spp. were recorded in the open vegetation and are key resources during the dry season. The availability of these flowering resources is, however, ephemeral which means they might not provide relief to hummingbirds should they adjust their diet preferences to plant species more likely to be present in open vegetation, modified areas

